# Angie-LAMP for diagnosis of human eosinophilic meningitis using dog as proxy: A LAMP assay for *Angiostrongylus cantonensis* DNA in cerebrospinal fluid

**DOI:** 10.1101/2022.12.19.521150

**Authors:** Vojtech Baláž, Phoebe Rivory, Douglas Hayward, Susan Jaensch, Richard Malik, Rogan Lee, David Modrý, Jan Šlapeta

## Abstract

**Background:** *Angiostrongylus cantonensis* (rat lungworm) is recognised as the leading cause of human eosinophilic meningitis, a serious condition observed when nematode larvae migrate through the CNS. Canine Neural Angiostrongyliasis (CNA) is the analogous disease in dogs. Both humans and dogs are accidental hosts, and rapid diagnosis is warranted. A highly sensitive PCR based assay is available but often not readily accessible in many jurisdictions. An alternative DNA amplification assay that would further improve the accessibility is needed. This study aimed to assess the diagnostic utility of a newly designed LAMP assay to detect DNA of globally distributed and invasive *A. cantonensis* and *Angiostrongylus mackerrasae*, the only other neurotropic *Angiostrongylus* species, which is native to Australia.

**Methodology/Principal Findings:** Cerebrospinal fluid (CSF) from dogs (2020-2022) with a presumptive diagnosis of *A. cantonensis* infection were received for confirmatory laboratory testing and processed for DNA isolation and ultrasensitive *Angiostrongylus* qPCR. A newly designed LAMP assay targeting AcanR3390 in a diagnostic laboratory setting was directly compared to the reference ultrasensitive qPCR for determination of presence of *A. cantonensis* DNA to aid the diagnosis of CNA. The LAMP assay (Angie-LAMP) allowed the sensitive detection of *A. cantonensis* DNA from archived DNA specimens (Kappa=0.81, 95%CI 0.69-0.92; *n*=93) and rapid single-step lysis of archived CSF samples (Kappa=0.77, 95%CI 0.59-0.94; *n*=52). Only *A. cantonensis* DNA was detected in canine CSF samples, and co-infection with *A. mackerrasae* using amplicon deep sequencing (ITS-2 rDNA) was not demonstrated. Both SYD.1 and AC13 haplotypes were detected using sequencing of partial *cox*1.

**Conclusions/Significance:** The Angie-LAMP assay is a useful molecular tool for detecting *Angiostrongylus* DNA in CSF of dogs and performs comparably to laboratory *Angiostrongylus* qPCR. Adaptation of single-step sample lysis improved potential applicability for effective diagnosis of angiostrongyliasis in a clinical setting for dogs and by extension for humans.

**Authors summary:** A potentially fatal disease, neural angiostrongyliasis, is caused by the rat lungworm (*Angiostrongylus cantonensis*). The parasite migrates into the spinal cord and brain of accidental hosts, such as humans and dogs, after ingestion of infective larvae. Recently, an ultrasensitive molecular assay which can detect tiny fragments of the parasite’s DNA was developed and has been used for confirmatory diagnosis. Although this assay outperforms previously developed assays, it requires clean DNA with specialised equipment in a laboratory setting. There is an urgent need for an alternative diagnostic method which is sensitive and portable, for deployment in the field and in the hospitals in remote areas or in low-income countries. The authors developed a fast and portable loop-mediated isothermal amplification (LAMP) assay that compares favourably to the ultra-sensitive PCR assay when tested using cerebrospinal fluid from dogs on the Australian east coast with presumptive neural angiostrongyliasis. Considering a ‘One Health’ approach to diagnostics, this assay enables portable emergency diagnostics equally suitable to humans, dogs and wildlife. The newly developed assay will also enable water supplies to be screened, as well as crustaceans and molluscs used as potential food sources, for presence of the parasite.

## Introduction

The ‘rat lungworm’ (*Angiostrongylus cantonensis*) is a nematode with a unique affinity for the central nervous system (CNS) found throughout the Asia-Pacific region, and more recently in the United States and Europe [1-8]. Currently, *A. cantonensis* is recognised globally as the leading cause of eosinophilic meningitis in humans, which is a serious condition observed when larvae migrate to the CNS [5, 9, 10]. It has been recorded as an etiological agent for meningitis in humans in at least 30 countries, including Australia, making it a parasite of increasing importance [11]. Humans are an accidental host when they deliberately or inadvertently ingest infected intermediate or paratenic hosts [9]. The rat lungworm traditionally cycles between the intermediate host – molluscs, and rats where it completes the life cycle and reaches the pulmonary arteries via an obligatory migration through the rat’s brain [12]. The infective larvae are generalists and will rapidly invade the CNS of many animals including primates, marsupials, bats, horses, dogs and birds [13-16]. The overt host response in the CNS leads to the associated neurological signs and symptoms, as well as severe inflammation of the meninges and brain [17]. Canine Neural Angiostrongyliasis (CNA) is a condition in dogs analogous to the human disease eosinophilic meningitis [18]. CNA manifests as severe neurological dysfunction, with signs such as hindlimb paresis, muscle wasting and urinary incontinence being common, characteristically associated with marked hyperaesthesia [19]. Naturally occurring cases of CNA are reported throughout most of coastal Queensland and eastern New South Wales in Australia [18-20]. There appears to be a “peak season” of CNA, with most cases occurring after unusually wet weather between April and June in QLD [18]. A similar pattern occurs in NSW where a spike in cases occurs around May [19]. Infection disproportionately affects younger dogs, speculated to be due to their higher propensity to eat slugs and snails out of curiosity, and possibly their naive immunity to nematodes [19, 20].

Diagnostics remain challenging in both humans and dogs due to the rapid onset of the clinical symptoms and signs, respectively [20-24]. The laboratory diagnostics rely on application of DNA amplification tests, with the recent introduction of ultrasensitive AcanR3390 real-time PCR assay demonstrated to be superior to previously used PCR and qPCR-based assays targeting rDNA in both humans and dogs [23, 25]. In an effort to further improve the accessibility of this assay it was adopted for Recombinase Polymerase Assay (RPA) that has the potential to become a point-of-contact molecular assay [25, 26]. As a compromise to qPCR and RPA, the loop-mediated isothermal DNA amplification (LAMP) using dsDNA binding fluorescent dye that can provide fast amplification, approximate quantification and product identity control, thus providing both speed and reliability [27, 28]. Such assays can be performed either in qPCR thermocycler or a portable device with appropriate fluorescence detection system [29].

This study aimed to assess the diagnostic utility of a newly designed LAMP assay targeting AcanR3390 in a diagnostic laboratory setting in a direct comparison with the reference ultrasensitive qPCR for determination of presence of *A. cantonensis* infection as a supportive tool to in the diagnosis of CNA. We used archived (2020-2022) cerebrospinal fluid (CSF) and DNA isolated from canine CSF from patients living along the east coast of Australia. The LAMP developed in this work (Angie-LAMP) was shown to be both sensitive and specific at detecting DNA from *A. cantonensis*; and the only other neurotropic *Angiostrongylus* species that is native to Australia, *A. mackerrasae*. The potential diagnostic applicability of the Angie-LAMP extends to other species and material, including human CSF from suspect neural angiostrongyliasis cases.

## Methods

### Sample collection

Cerebrospinal fluid (CSF) from 111 domestic dogs were provided by Vetnostics (Laverty Pathology - North Ryde Laboratory, Sydney) and Veterinary Pathology Diagnostics Services (University of Sydney). In total, 108 samples were submitted for confirmatory molecular diagnostics of *A. cantonensis* infections, with signalment suggesting infectious cause and clinician’s suspicion for CNA (S1 Table). The remaining three CSF samples did not have clinical suspicion of CNA (Ag70, Ag71, Ag107) (S1 Table). Material (CSF) was collected by registered veterinary practitioners in accordance with the Veterinary Practice and Animal Welfare Acts. Material was provided for the purpose of diagnostics and results used in decision-making by the veterinary practitioner with residual material made available for this work. Results and material used here were deidentified.

### DNA isolation from cerebrospinal fluid (CSF)

Genomic DNA was purified from 100μL of CSF samples using the Monarch® Genomic DNA Purification Kit (New England Biolabs, Australia) according to the manufacturer’s instructions, using the Mammalian Whole Blood protocol. Up to 100μL of CSF was lysed, with insufficient samples being topped up to the required volume with phosphate-buffered saline (PBS). Each batch of samples processed included a ‘blank’ extraction control. Samples were eluted with 75μL of pre-heated gDNA Elution Buffer, and the final centrifugation step was repeated for maximum gDNA yield. DNA and remaining unused CSF were stored at −20°C prior to further processing.

### Ultrasensitive AcanR3390 *Angiostrongylus* qPCR

A probe-based qPCR assay targeting a repeat sequence on contig 3990 (AcanR3390) of *A. cantonensis* described by Sears et al. [25] was locally optimised and used as a veterinary diagnostic assay at the Laboratory of Veterinary Parasitology, University of Sydney. Duplicate reactions were run at a final volume of 20μL, including 2μL of gDNA and 10μL Luna® Universal Probe qPCR Master Mix (New England Biolabs, Australia). Final concentrations of the forward (S0947) and reverse (S0948) primers were 0.4μM, and FAM probe (S0949) at a final concentration of 0.1μM. Primers and probes were sourced from Integrated DNA Technologies, Inc. (Australia). All qPCR runs included a ‘blank’ extraction control, ddH_2_O no-template control, and positive *A. cantonensis* control for the detection of contamination and assurance of qPCR run success. qPCRs were performed in a CFX96 Touch Real-Time PCR Detection System (Bio-Rad Laboratories, Inc.) with the following cycling conditions: 95°C for 3 minutes; and 40 cycles of 95°C for 5 seconds, and 60°C for 15 seconds. Amplification curves and cycle threshold (C_t_)-values were recorded using CFX Maestro Software 2.3 (Bio-Rad Laboratories, Inc.).

A region of the mammalian glyceraldehyde 3-phosphate dehydrogenase (G3PDH) gene was targeted by a separate probe-based qPCR assay, which was run in parallel with the AcanR3390 assay to ensure that DNA was successfully isolated. An additional positive control containing DNA isolated from whole dog blood was included in these runs. Reagent volumes and concentrations were the same as outlined above, but instead using G3PDH forward and reverse primers (S1072 and S1073, respectively), and Cy5 probe (S1074) described by Peters et al. [30].

Currently used interpretation workflow for ultrasensitive AcanR3390 *Angiostrongylus* qPCR utilised by the Sydney School of Veterinary Science is as follows. Resulting C_t_ values from duplicate qPCR of the ultrasensitive *Angiostrongylus* DNA alongside C_t_ values from qPCR for presence of mammalian DNA are recorded. Sample/s are run alongside the isolated ‘blank’ extraction control sample. (1) Mammalian DNA qPCR is positive (C_t_ < 40) for the sample DNA, while the ‘blank’ is negative for both mammalian DNA and *Angiostrongylus* DNA (C_t_ ≥ 40). All samples in this set passed this step. (2) If duplicate *Angiostrongylus* DNA qPCR returns C_t_ ≤ 35 in both qPCRs, the sample is considered ‘strong positive’ for *Angiostrongylus* DNA. (3) If one of the qPCRs returns C_t_ ≤ 35 and the other C_t_ > 35 (including C_t_ ≥ 40), the sample is considered ‘positive’ for *Angiostrongylus* DNA. (4) If duplicate *Angiostrongylus* DNA qPCR returns C_t_ ≤ 38 but C_t_ > 35 in both qPCRs, the sample is considered ‘weak positive’ for *Angiostrongylus* DNA. (5) The sample is considered ‘equivocal’ for *Angiostrongylus* DNA if one C_t_ > 38 and the other does not amplify (return C_t_ ≥ 40), or one is C_t_ > 35 but the other does not amplify, or both C_t_ > 38. (6) If both qPCRs do not amplify (return C_t_ ≥ 40), the sample is considered ‘negative’ for *Angiostrongylus* DNA.

### Molecular determination of *Angiostrongylus* species in CSF

We adopted an ITS-2 rDNA qPCR diagnostic assay for amplicon metabarcoding and Next-Generation Sequencing (NGS) to allow us to determine *Angiostrongylus* species. The qPCR primers adapted from Fang et al. [31] target a ∼140bp region of ITS-2 rDNA; [S1060] AngioITS2_FOR (*TCG TCG GCA GCG TCA GAT GTG TAT AAG AGA CAG* CCA GTT TTG GTG AAG AAT AA) and [S1061] AngioITS2_REV (*GTC TCG TGG GCT CGG AGA TGT GTA TAA GAG ACA G*AC ACG ACG GTA ACA ATG ACA), nucleotides in italic represent the Nextera (Illumina) adapter. This region includes a single nucleotide polymorphism (SNP) (G-to-A) at the 83^rd^ position which discriminates *A. cantonensis* from *A. mackerrasae* [32]. A selection of individual dog DNA CSF samples (*n*=35), and DNA from voucher *A. cantonensis* (*n*=6) and *A. mackerrasae* (*n*=2) specimens from Valentyne et al. [33] were subjected to barcoding amplification and processing for NGS at the Ramaciotti Centre for Genomics, University of New South Wales, Australia using Illumina MiSeq v2 250 PE. Obtained FastQ files were processed using a local DADA2 pipeline and curated database of reference sequences via CLC Main Workbench v22 (Qiagen, CLC bio). The output (in the form of ASV counts per sample) was manually analysed in Excel to identify samples with >200 reads matching a reference *Angiostrongylus* sequence. FastQ files were deposited in Sequence Read Archive (SRA) under the BioProject: PRJNA912228.

To determine the *cox*1 haplotype of *A. cantonensis*, a partial *cox*1 was amplified and sequenced according to Mallaiyaraj Mahalingam et al. [32]. Briefly, primers AngiCOI_forward (S0963) and AC1R (S0966) were used in a qPCR reaction with Luna® Universal qPCR Mastermix (New England Biolabs, Australia). The qPCR reactions were run on the CFX96 Touch Real-Time PCR Detection System (BioRad, Australia) and analysed using the corresponding CFX Maestro 2.3 software (BioRad, Australia). Cycling conditions were as follows: 95 °C for 60 s, followed by 40 cycles at 95 °C for 15 s and 55 °C for 30 s. Results were considered positive if the melt curve profile corresponded to that of *A. cantonensis* control and reactions submitted for bidirectional sequencing at Macrogen Inc. (Seoul, Korea). The DNA chromatographs were visually inspected in CLC Main Workbench v22 (Qiagen, CLC bio) and matched against the reference alignment of all known *cox1* haplotype sequences from GenBank. Newly obtained *cox*1 were deposited in GenBank (OQ029497-OQ029501). Additional associated files are available from LabArchives (https://dx.doi.org/10.25833/z7zq-9e13).

### Loop-mediated isothermal DNA amplification for detection of *Angiostrongylus* DNA - Angie-LAMP assay

Primers for Angie-LAMP were designed using the GLAPD online tool [34] using the published Acan3990 tandem repeats sequence as a template [25]. The selected primers and optimal concentrations used are presented in Table 1. All Angie-LAMP reactions (25 μL) were run using Isothermal Master Mix (ISO-004, Optigene, UK) according to manufacturer’s instructions in a portable machine Genie® II and/or Genie® III (Optigene, UK) equipped for reading, recording and visualisation of the FAM fluorescence.

**Table 1:**
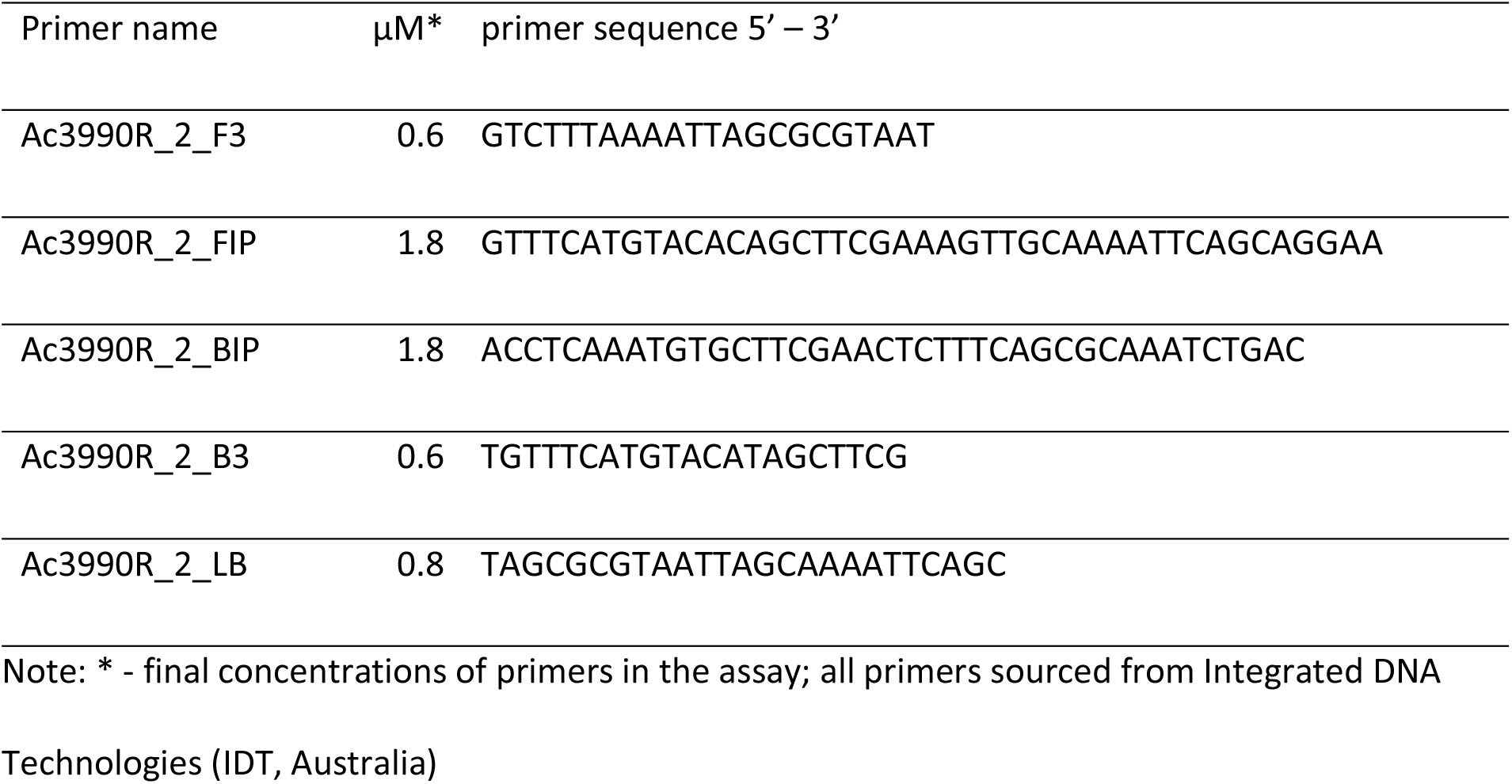
Angie-LAMP assay primers and primer concentrations.

The Angie-LAMP assay was tested against DNA extracted from various lineages of *A. cantonensis* (Fatu Hiva lineage FH.1, lineage in Sydney SYD.1, and Thai lineage in Sydney AC13), and other *Angiostrongylus* species (*A. mackerrasae, A. chabaudi, A. costaricensis, A. daskalovi* and *A. vasorum*). In addition, relevant DNA from hosts (human, dog) and DNA from migratory parasites that can be present in CSF of dogs (*Toxocara, Ophidiascaris, Ancylostoma, Uncinaria*) were also tested. The limit of detection was calculated by testing 10-fold serial dilutions of DNA from *A. cantonensis* L1 larvae.

Two *Angiostrongylus* DNA positive CSF samples (Ag66, Ag77) were initially used to evaluate a ‘point-of care’ applicability of the Angie-LAMP. We tested (i) neat CSF sample, (ii) CSF sample heated at 100°C for 10 minutes, and (iii) CSF sample treated with PrepMan™ Ultra (Thermo-Fisher Scientific) and heated at 100°C for 10 minutes. The selected samples were analysed at different dilutions (undiluted to 1:1,000) and used immediately for Angie-LAMP.

Archived DNA isolated from dog CSF samples (*n*=111) and available neat CSF samples from a subset of the DNA samples for which retained CSF was available (*n*=57) was used to compare the standard laboratory diagnostic procedure (DNA isolated using extraction kit, followed by *Angiostrongylus* qPCR). Neat CSF samples (10 μL) were mixed with equal volume PrepMan™ Ultra (Thermo-Fisher Scientific) reagent in 0.2 mL PCR 8-well strips and heated to 100 °C for 10 min (T100; BioRad, Australia) and used immediately. Angie-LAMP reactions included either 5 μL of DNA isolated from CSF or 5 μL of the CSF treated with PrepMan™ Ultra reagent and run in Genie III (OptiGene, UK) with 30 minutes amplification at 65°C and subsequent anneal temperature analysis (98°C to 80°C). Negative (ddH_2_O) and positive (*A. cantonensis* SYD.1) DNA controls were included. Two values obtained for each tested sample were the time of amplification detection in minutes (in principle comparable to C_t_ in qPCR) and annealing temperature (when primers attach to the DNA product and so cause measurable change in fluorescence of the dsDNA intercalating dye). The processing of raw data in Genie Explorer software (OptiGene, UK) included control of fluorescence change during amplification and annealing analysis (shape, angle of curves). Time of detected amplification calculation had to be adjusted in some runs to eliminate erroneous readings caused by background changes in fluorescence (smoothing by 2-3 points, detection threshold 0.010-0.008 ratio). Anneal derivative analysis (smoothing 2-5 points, peak detection full, threshold varied from 4,000-5,000 derivative). If amplification read more values due to atypical shape of the amplification derivative, the correct value was obtained manually in Genie Explorer software, reading the peak value. A sample was considered ‘positive’ if amplification was detected before 25^th^ minute and quantified anneal temperature was within 0.5 °C from the positive controls. If only single reading of either amplification time or anneal temperature, or late amplification occurred (>25 minutes), we interpreted the result as ‘equivocal’. No amplification within 30 minutes or amplification of a product with anneal temperature outside the set range was considered as a ‘negative’ LAMP result.

### Data analysis

Data were imported and analysed in GraphPad Prism 9.4.1 (GraphPad Software, US). Data were visualised as scatterplots and histograms, analysed using Bland-Altman plot and linear regression to find the best line that predicts Y from X and interpreted using contingency tables. Means were compared using *t*-tests. Data were tested for normality (D’Agostino & Pearson test), 95% confidence intervals calculated, and significance was set at 0.05. Kappa was interpreted using the scale of Landis and Koch [35] and Bland Altman plot according to Giavarina [36].

## Results

### Performance of AcanR3390 and Angie-LAMP assays

The CSF samples included in this study (*n*=111) were processed for DNA isolation followed by the diagnostic workflow for ultrasensitive *Angiostrongylus* qPCR (Figure 1A; Table 2). In total, 64/111 (58%) samples were positive, comprising 53/111 (48%) strong positives, 5/111 (5%) positives and 6/111 (5%) weak positives. Nine samples were considered equivocal for presence of *Angiostrongylus* DNA, while 38/111 (34%) were qPCR-negative for *Angiostrongylus* DNA.

**Table 2:** Summary of molecular diagnostics for the detection of *Angiostrongylus* DNA.

| Angie-LAMP (DNA) / Angie-LAMP (CSF) |  |  |  |  |
| --- | --- | --- | --- | --- |
| *Angiostrongylus* qPCR | Positive | Equivocal | Negative | Total |
| <sup>a</sup> Strong positive | 47/27 | 0/2 | 6/3 | 53/32 |
| <sup>b</sup> Positive | 1/1 | 1/0 | 3/2 | 5/3 |
| <sup>c</sup> Weak positive | 0/0 | 2/0 | 4/0 | 6/0 |
| <sup>d</sup> Equivocal | 0/0 | 0/0 | 9/1 | 9/1 |
| <sup>e</sup> Negative | 0/1 | 2/2 | 36/18 | 38/21 |
| Total | 48/29 | 5/4 | 58/24 | 111/57 |

Note:
<sup>a</sup> $C_t \leq 35$ in both
<sup>b</sup> one $C_t \leq 35$ and other $C_t > 35$ including $C_t \geq 40$ (no amplification)
<sup>c</sup> both $C_t \leq 38$ but $C_t > 35$
<sup>d</sup> one $C_t > 38$ and other $C_t \geq 40$ (no amplification), or one $C_t > 35$ but other $C_t < 40$ , or both $C_t > 38$
<sup>e</sup> $C_t \geq 40$ in both (no amplification)

**Figure 1.**
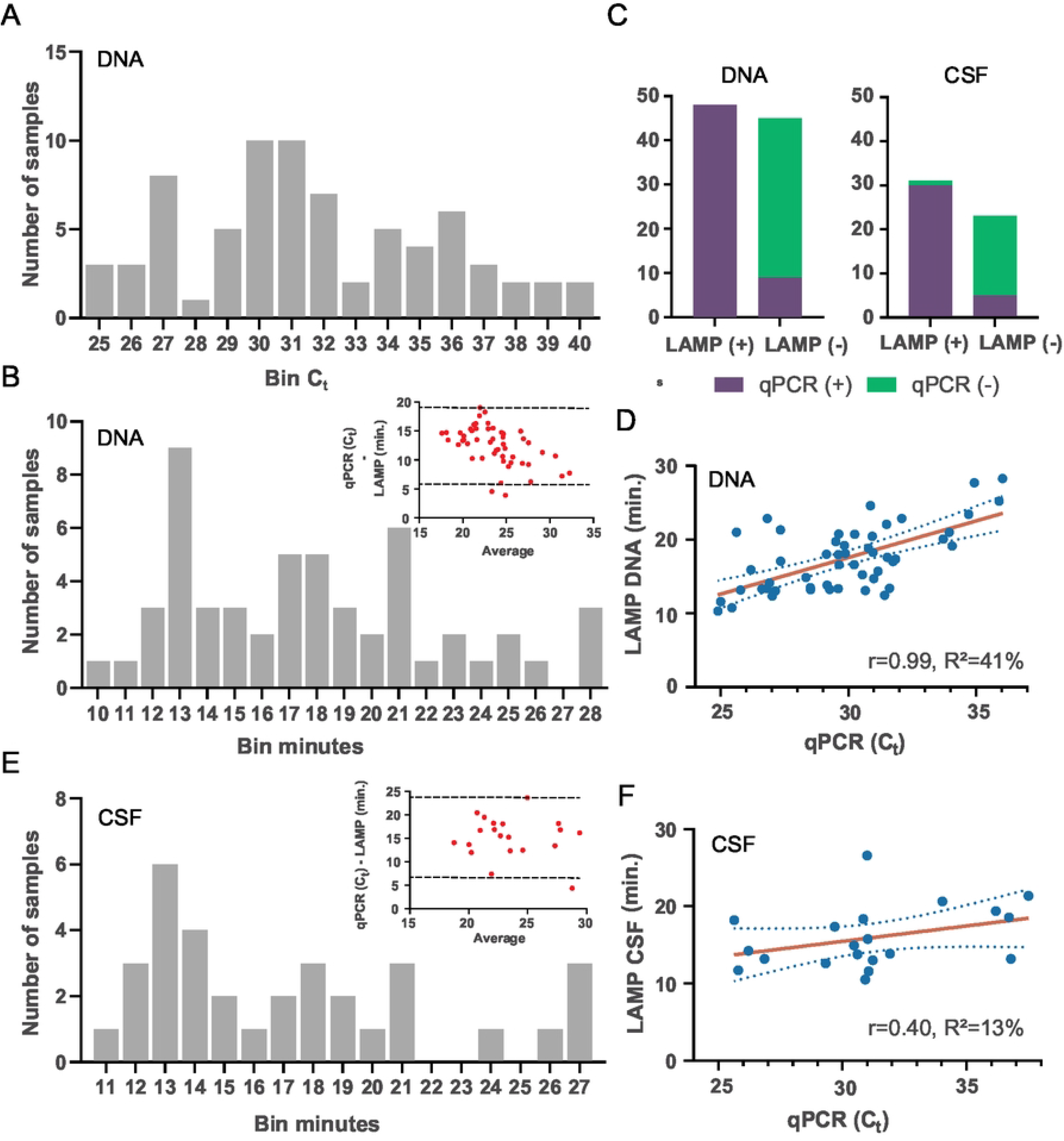
Performance of Angie-LAMP assay targeting AcanR3390 with direct comparison to a reference ultrasensitive qPCR to detect *Angiostrongylus* DNA in canine cerebrospinal fluid (CSF). (A) Histograms of the distribution of the qPCR C_t_ values of *Angiostrongylus* positive DNA samples (C_t_<40, *n*=73). DNA was isolated from canine CSF samples and qPCR assay was run in duplicate. Avg. C_t_ value was used for the histogram. Each ‘Bin C_t_’ represents the number of samples within that category, for example ‘40’ = C_t_ values from 39 to <40. (B) Distribution of Angie-LAMP amplification time (‘minutes’) when DNA isolated from canine CSF returned positive signal (*n*=53). Individual positive values were used for the histogram. Each ‘Bin minutes’ as described above. Inset: Bland Altman plot with 95% confidence agreement (dotted line). (C) Stacked bar plots of the values obtained from a contingency table comparing the sensitivity and specificity of the Angie-LAMP assay to the reference ultrasensitive qPCR assay. (D) Scatterplot with linear regression line plotted and 95% confidence interval (dotted line) shown for DNA run using Angie-LAMP [LAMP DNA min.] (R^2^, and slope ‘r’ is shown). (E) Distribution of Angie-LAMP amplification time (‘minutes’) for available boiled CSF mixed with PrepMan™ Ultra reagent that returned positive signal (*n*=33). Individual positive values were used for the histogram. Each ‘Bin minutes’ as described above. Inset: Bland Altman plot with 95% confidence agreement (dotted line). (F) Scatterplot with linear regression line plotted and 95% confidence interval (dotted line) shown for available boiled CSF mixed with PrepMan™ Ultra reagent using Angie-LAMP [LAMP CSF min.] (R^2^, and slope ‘r’ is shown).

All 111 DNA samples isolated from CSF were subject to Angie-LAMP, 48/111 (43%) were positive 58/111 (52%) were negative and 5/111 (5%) returned an equivocal amplification result (Figure 1B; Table 2). Weak positive qPCR samples were either Angie-LAMP negative (4/6) or Angie-LAMP equivocal (2/6). All nine qPCR equivocal samples returned negative result using Angie-LAMP. In addition, three samples using Angie-LAMP returned equivocal result (one qPCR positive, two qPCR negative samples). All equivocal and qPCR weak positive results were not considered further. Using 93 paired samples, the agreement (Kappa = 0.81; 95%CI 0.69 to 0.92) between the two tests was considered ‘almost perfect agreement’ to ‘substantial agreement’ using the scale of Landis and Koch [35]. Bland-Altman bias was 12.4 (SD=3.4) with 95% limit of agreement 5.8 to 19 (Figure 1B - inset). Assuming the qPCR as the gold standard reference test, the Angie-LAMP sensitivity is 84.2% (95%CI 72.1% to 92.5%) and specificity is 100% (95%CI 90.3% to 100%) (Figure 1C). Linear regression demonstrated that the slope between qPCR and Angie-LAMP was significantly non-zero (F=33.93, DFn=1, DFd=49, P-value<0.0001), with a moderate coefficient of determination (R^2^=41%) (Figure 1D).

To further consider the diagnostic utility of Angie-LAMP, we attempted to bypass DNA extraction as a separate step and utilised a single step protocol for cell lysis with PrepMan™ Ultra reagent. We assessed 57 canine CSF samples previously used for DNA isolation and tested for presence of *Angiostrongylus* DNA (32 strong positive, 3 positive, 21 negative, 1 equivocal) (Figure 1E; Table 2). The agreement of the Angio-LAMP on CSF samples treated with PrepMan™ Ultra with *Angiostrongylus* qPCR showed ‘substantial agreement’ (Kappa = 0.77; 95%CI 0.59 to 0.94) using the scale of Landis and Koch [35]. Bland-Altman bias was 15 (SD=4.4) with 95% limit of agreement 6.7 to 24 (Figure 1E inset). Using the qPCR as the reference test, the Angie-LAMP assay using CSF treated with PrepMan™ Ultra reagent had a sensitivity of 85.7% (95%CI 69.7% to 95.2%) and specificity of 94.7% (95%CI 74.0% to 99.9%) (Figure 1C). Linear regression demonstrated that the slope between qPCR and Angie-LAMP on CSF treated with PrepMan™ Ultra was not significantly ‘non-zero’ (F=2.59, DFn=1, DFd=18, P-value = 0.13; R^2^=13%) (Figure 1F).

The use of PrepMan™ Ultra reagent was initially evaluated using serial dilutions of two CSF samples with *Angiostrongylus* DNA (*Angiostrongylus* qPCR: Ag66, avg. C_t_ value = 27.0; Ag77, avg. C_t_ value = 27.0). We tested (i) neat CSF, (ii) boiled CSF and (iii) boiled CSF mixed with PrepMan™ Ultra reagent in the Angie-LAMP. Neat, undiluted samples returned positive, but late Angie-LAMP signal (>25 minutes); further dilution yielded no amplification. Boiled samples (undiluted to 1:1,000) returned inconsistent annealing profiles that did not match the positive control (DNA). Both boiled CSF samples mixed with PrepMan™ Ultra reagent (undiluted to 1:1,000) returned expected positive Angie-LAMP signal and product that matched the annealing profile of the positive control (DNA of *A. cantonensis*).

### Confirmed presence of *Angiostrongylus cantonensis* in CSF of dogs

*Angiostrongylus* species identification was successful in 40% (14/35) of samples that were processed for ITS-2 rDNA deep sequence analysis. Of the 64 *Angiostrongylus* AcanR3990-positive individual dog CSF samples, 35 were processed using the ITS-2 rDNA amplification. Overall, 66% (23/35) produced ITS-2 rDNA amplicon with primers adopted for deep sequencing. Fourteen ITS-2 rDNA amplicons produced sequence reads that matched *Angiostrongylus* ITS-2 rDNA. All (14/14) of analysed dog CSF samples were identified unequivocally as containing only *A. cantonensis* DNA. The mean C_t_ value of the successfully identified DNA was ∼8 cycles lower than those that were unsuccessful (paired *t*-test, P value < 0.001). Amplification and deep sequencing of voucher DNA of *A. cantonensis* and *A. mackerrasae* confirmed the G-to-A SNP within ITS-2 rDNA.

For 33 *Angiostrongylus* qPCR positive samples, we attempted to PCR amplify partial *cox*1 to enable identification of *Angiostrongylus* species and mtDNA haplotype. Five/33 (15%) produced DNA sequencing results; four matched perfectly to the SYD.1 haplotype (Ag8, Ag11, Ag13, Ag72) and one perfectly matched AC13 haplotype (Ag6).

## Discussion

The recently developed ultrasensitive qPCR targeting the AcanR3390 repetitive element can detect *A. cantonensis* in samples with as little as 0.00001 of a single third stage larva [25]. Such performance improved the ability to diagnose rat lungworm disease and outperforms previously used assays in clinically confirmed case of CNA [23]. While the assay is highly sensitive it requires collection of CSF and DNA isolation in a specialised laboratory setting before the assay can be performed. Our approach adopted LAMP technology to target the respective element, because LAMP can be used in field- or point of care conditions and works using various samples, i.e. CSF. The sensitivity performance of LAMP is consistent with qPCR. Analogous to previous studies targeting AcanR3390, Angie-LAMP is highly specific for globally emerging *A. cantonensis* and so does not cross react with any other viruses, bacteria, protozoa and helminths, except *A. mackerrasae* [25, 26].

A closely related sibling species native to Australia, *A. mackerrasae*, shares a life cycle and neurotropic tendencies to that of *A. cantonensis*; however, *A. cantonensis* is universally assumed to be the causative agent of human eosinophilic meningitis and CNA [21, 37]. Although, it is speculated that *A. mackerrasae* could cause CNA – especially as a patent infection was discovered in a black flying fox (*Pteropus alecto*) in 2013 [38, 39]. In this study using a deep sequencing approach and recent canine CSF samples (2020-2022), we demonstrate an absence of *A. mackerrasae* DNA which supports the dogma that *A. cantonensis* is the sole culprit in CNA cases [32]. Unlike the previous study by Mallaiyaraj Mahalingam et al. [32], we adopted the ITS-2 assay for deep amplicon sequencing to detect potential co-infections between the two species. The absence of *A. mackerrasae* DNA in dogs with CNA in the current study may result from *A. mackerrasae*’s innate preference for native rat species such as *Rattus fuscipes* (bush rat) and *R. lutreolus* (Australian swamp rat) due to undergone co-evolution [37]. Bush and swamp rats are known to avoid dispersal through developed areas where invasive *Rattus* spp. thrive, which may reduce cross-over of domestic dogs and native rats [40-42]. Alternatively, *A. mackerrasae* is simply a specialist not capable of accidently infecting dogs.

Globally, many *cox*1 haplotypes of *A. cantonensis* have been characterised [33, 43]. The analysis of the partial *cox1* region from dog CSF in our study confirms that Australian dogs currently host at least two known haplotypes; the established SYD.1, and the more recently recognised AC13 from Thailand [32, 44]. The presence of additional haplotypes locally is not yet known, but is speculated due to the ongoing invasion of *A. cantonensis* strains into previously non-endemic countries [45, 46]. Experimental infection of rats with various Taiwanese (H and P strains) and Brazilian (ac8 and ac9) *A. cantonensis* haplotypes demonstrated differences in infectivity and fecundity [47, 48]. Further work should aim to determine if the two now endemic Australian haplotypes have different virulence properties, geographical ranges, host preferences or propensity to infect non-permissive hosts, such as dogs, tawny frog mouths, bats and humans.

The newly designed LAMP assay targeting AcanR3390 demonstrated substantial agreement without the need to extract purified DNA, and thus diagnostic utility under laboratory conditions. Both the correlation and the Bland-Altman plot suggest poor agreement between the quantitative readings from qPCR (C_t_ values) and Angie LAMP (minutes) tests, nevertheless the qualitative observations provide robust Kappa agreement outcome that is clinically relevant. One, however, has to consider if LAMP is a useful alternative diagnostic modality for clinical scenarios in both veterinary and medical settings. Infections with *A. cantonensis* are medical as well as veterinary emergencies that requires rapid intervention to alleviate the overt host response in the meninges as well as administration of antiparasiticides [24, 49]. Clinical signalment and signs associated with the syndrome often lead to presumptive diagnosis and initiation of treatment regardless of early laboratory data, as CSF collection can be financially prohibitive or not feasible due to the locations where these patients are situated. In situations where CSF is collected, the typical finding is eosinophilic pleocytosis, or an increased percentage of eosinophils [22, 50]. Diagnostic PCR, including the ultrasensitive AcanR3390, are not ‘rule out’ tests, because in clinically confirmed cases of CNA summarised by Lee et al. [23] only 81% (48/59) of affected dogs tested positive for presence of *Angiostrongylus* DNA, the remainder being diagnosed on the basis of detecting specific antibodies directed against *A. cantonensis* in CSF using ELISA. In veterinary situation, DNA tests often take 1-2 days and no private veterinary hospitals have molecular diagnostic capacity on site. Availability of the portable instrument utilised in this study and the ease of preparations of the samples without the need for DNA extraction as a separate procedure can enable on site diagnostics for moderate to large referral hospitals. The turnaround from collecting the CSF to receiving the result can be achieved in under 1 hour (CSF preparation 10 minutes, LAMP 30 minutes, 10 minutes sample and reagents manipulation). In fact, our results show that utilising LAMP will only slightly increase the false negative rate compared to reference laboratory results.

## Conclusion

We have developed new Angie-LAMP that performs favourably on CSF of dogs if compared to referral diagnostics laboratory qPCR method following DNA isolation. Replacing the DNA isolation using single-step preparation of CSF is feasible in current clinical settings and thus Angie-LAMP can enable ‘point-of-care’ diagnostics using CSF. One of the main reasons of this study was to consider a ‘One Health’ approach to diagnostics, because the serious emergency consequences of this parasite on both humans and dogs. Our archival material of CSF samples from 2020-2022, where CNA was presumed, potentially represents one of the largest sample sizes of CSF specimens to evaluate new test such as Angie-LAMP. While CNA is frequently recorded in eastern Australia [23], the equivalent human condition eosinophilic meningitis is rare and sporadic, with far fewer detected cases and access to archived CSF collections. The pathogenesis of the disease in dogs and humans enables us to consider the performance of Angie-LAMP on canine CSF as proxy to performance on human CSF. In both species, the CSF has abnormally high number of cells that are mostly eosinophils. The assay does not amplify human or canine DNA, nor does it amplify other potential neuropathogens including parasites that could be present in CSF. Our results show that Angie-LAMP is a robust tool suitable for use in either canine or human diagnostics. Furthermore, Angie-LAMP is ideally suited for testing in the field for screening water supplies, food (crustaceans and molluscs as food source), snail infested waters (eDNA) or rat faecal pellets for presence of *Angiostrongylus*. As far as veterinary and human diagnostics, the retrieval of CSF via spinal tap is an invasive and potentially traumatic procedure. Evaluation of the validated Angie-LAMP assay on other more accessible diagnostic material (such as well-timed paired whole blood, sera and urine specimens) is urgently needed.

## Acknowledgements

This work was supported by the University of Sydney Research Training Program (RTP) Scholarship (Australian Government) and RTP Tuition Fee Offset (University of Sydney) to PR; development of the LAMP assay was funded by SEAEUROPEJFS19IN-053 (ASEAN-EU Cooperation in Science, Technology and Innovation, Southeast Asia - Europe Joint Funding Scheme 2019). The funders had no role in the design and conduct of the study, nor the decision to prepare and submit the manuscript for publication. We thank GeneWorks (Australia) for lending us Genie® III (Optigene, UK).

## Author Contributions

*Vojtech Baláž*: Conceptualization, Data Curation, Formal Analysis, Investigation, Methodology, Funding Acquisition, Resources, Writing – Original Draft Preparation, Writing – Review & Editing

*Phoebe Rivory*: Conceptualization, Data Curation, Formal Analysis, Investigation, Methodology, Writing – Original Draft Preparation, Writing – Review & Editing

*Douglas Hayward*: Resources, Writing – Review & Editing

*Susan Jaensch*: Resources, Writing – Review & Editing

*Richard Malik*: Writing – Review & Editing

*Rogan Lee*: Resources, Writing – Review & Editing

*David Modry*: Conceptualization, Funding Acquisition, Resources, Writing – Review & Editing

*Jan Šlapeta*: Conceptualization, Data Curation, Formal Analysis, Funding Acquisition, Resources, Supervision, Writing – Original Draft Preparation, Writing – Review & Editing

## Supplementary Material

**S1 Table:** Summary of canine cerebrospinal fluid samples used in this study with diagnostic results

## References

1. York EM, Creecy JP, Lord WD, Caire W. Geographic range expansion for rat lungworm in North America. Emerging Infect Dis. 2015;21(7):1234–6. doi: 10.3201/eid2107.141980.

2. Paredes-Esquivel C, Sola J, Delgado-Serra S, Puig Riera M, Negre N, Miranda MÁ, et al. Angiostrongylus cantonensis in North African hedgehogs as vertebrate hosts, Mallorca, Spain, October 2018. Euro Surveill. 2019;24(33):1900489. doi: 10.2807/1560-7917.ES.2019.24.33.1900489.

3. Delgado-Serra S, Sola J, Negre N, Paredes-Esquivel C. Angiostrongylus cantonensis nematode invasion pathway, Mallorca, Spain. Emerg Infect Dis. 2022;28(6):1163–9. doi: 10.3201/eid2806.212344.

4. Galán-Puchades MT, Gómez-Samblás M, Osuna A, Sáez-Durán S, Bueno-Marí R, Fuentes MV. Autochthonous Angiostrongylus cantonensis lungworms in urban rats, Valencia, Spain, 2021. Emerg Infect Dis. 2022;28(12):2564–7. doi: 10.3201/eid2812.220418.

5. Cowie RH, Ansdell V, Panosian Dunavan C, Rollins RL. Neuroangiostrongyliasis: global spread of an emerging tropical disease. Am J Trop Med Hyg. 2022. doi: 10.4269/ajtmh.22-0360.

6. Jarvi S, Prociv P. Angiostrongylus cantonensis and neuroangiostrongyliasis (rat lungworm disease): 2020. Parasitology. 2021;148(2):129–32. doi: 10.1017/S003118202000236X.

7. Gonzalvez M, Ruiz de Ybanez R. What do we know about Angiostrongylus cantonensis in Spain? Current knowledge and future perspectives in a globalized world. Transbound Emerg Dis. 2022;69(5):3115–20. doi: 10.1111/tbed.14393.

8. Turck HC, Fox MT, Cowie RH. Paratenic hosts of Angiostrongylus cantonensis and their relation to human neuroangiostrongyliasis globally. One Health. 2022;15:100426. doi: 10.1016/j.onehlt.2022.100426.

9. Cowie RH. Biology, systematics, life cycle, and distribution of Angiostrongylus cantonensis, the cause of rat lungworm disease. Hawaii J Med Public Health. 2013;72(6):6–9.

10. Barratt J, Chan D, Sandaradura I, Malik R, Spielman D, Lee R, et al. Angiostrongylus cantonensis: a review of its distribution, molecular biology and clinical significance as a human pathogen. Parasitology. 2016;143(9):1087–118. doi: 10.1017/S0031182016000652.

11. Kliks MM, Palumbo NE. Eosinophilic meningitis beyond the Pacific Basin: The global dispersal of a peridomestic zoonosis caused by Angiostrongylus cantonensis, the nematode lungworm of rats. Soc Sci Med. 1992;34(2):199–212. doi: 10.1016/0277-9536(92)90097-A.

12. Mackerras MJ, Sandars DF. The life history of the rat lung-worm, Angiostrongylus cantonensis (Chen) (Nematoda: Metastrongylidae). Aust J Zool. 1955;3(1):1–21. doi: 10.1071/ZO9550001.

13. Monks DJ, Carlisle MS, Carrigan M, Rose K, Spratt D, Gallagher A, et al. Angiostrongylus cantonensis as a cause of cerebrospinal disease in a yellow-tailed black cockatoo (Calyptorhynchus funereus) and two tawny frogmouths (Podargus strigoides). J Avian Med Surg. 2005;19(4):289–93. doi: 10.1647/2004-024.1.

14. Patial S, Delcambre BA, DiGeronimo PM, Conboy G, Vatta AF, Bauer R. Verminous meningoencephalomyelitis in a red kangaroo associated with Angiostrongylus cantonensis infection. J Vet Diagn. 2022;34(1):107–11. doi: 10.1177/10406387211037664.

15. Reddacliff LA, Bellamy TA, Hartley WJ. Angiostrongylus cantonensis infection in grey-headed fruit bats (Pteropus poliocephalus). Aust Vet J. 1999;77(7):466–8. doi: 10.1111/j.1751-0813.1999.tb12095.x.

16. Ma G, Dennis M, Rose K, Spratt D, Spielman D. Tawny frogmouths and brushtail possums as sentinels for Angiostrongylus cantonensis, the rat lungworm. Vet Parasitol. 2013;192(1-3):158–65. doi: 10.1016/j.vetpar.2012.11.009.

17. Martins YC, Tanowitz HB, Kazacos KR. Central nervous system manifestations of Angiostrongylus cantonensis infection. Acta Trop. 2015;141:46–53. doi: 10.1016/j.actatropica.2014.10.002.

18. Mason KV. Canine neural angiostrongylosis: the clinical and therapeutic features of 55 natural cases. Aust Vet J. 1987;64(7):201–3. doi: 10.1111/j.1751-0813.1987.tb15181.x.

19. Lunn J, Lee R, Smaller J, MacKay B, King T, Hunt G, et al. Twenty two cases of canine neural angiostronglyosis in eastern Australia (2002-2005) and a review of the literature. Parasit Vectors. 2012;5(1):70. doi: 10.1186/1756-3305-5-70.

20. Walker AG, Spielman D, Malik R, Graham K, Ralph E, Linton M, et al. Canine neural angiostrongylosis: a case-control study in Sydney dogs. Aust Vet J. 2015;93(8):264. doi: 10.1111/avj.12357.

21. Prociv P, Spratt DM, Carlisle MS. Neuro-angiostrongyliasis: unresolved issues. Int J Parasitol. 2000;30(12-13):1295–303. doi: 10.1016/s0020-7519(00)00133-8.

22. Wang Q-P, Lai D-H, Zhu X-Q, Chen X-G, Lun Z-R. Human angiostrongyliasis. Lancet Infect Dis. 2008;8(10):621–30. doi: 10.1016/S1473-3099(08)70229-9.

23. Lee R, Pai T-Y, Churcher R, Davies S, Braddock J, Linton M, et al. Further studies of neuroangiostrongyliasis (rat lungworm disease) in Australian dogs: 92 new cases (2010–2020) and results for a novel, highly sensitive qPCR assay. Parasitology. 2021;148(2):178–86. doi: 10.1017/S0031182020001572.

24. Prociv P. Need for critical rethinking in clinical approaches to neuroangiostrongyliasis. Am J Trop Med Hyg. 2019;101(4):951. doi: 10.4269/ajtmh.19-0546a.

25. Sears WJ, Qvarnstrom Y, Dahlstrom E, Snook K, Kaluna L, Baláž V, et al. AcanR3990 qPCR: A novel, highly sensitive, bioinformatically-informed assay to detect Angiostrongylus cantonensis infections. Clin Infect Dis. 2020;73:e1594–e600. doi: 10.1093/cid/ciaa1791.

26. Sears WJ, Qvarnstrom Y, Nutman TB. RPAcan3990: An ultrasensitive recombinase polymerase assay to detect Angiostrongylus cantonensis DNA. J Clin Microbiol. 2021;59(9):e0118521. doi: 10.1128/jcm.01185-21.

27. Notomi T, Okayama H, Masubuchi H, Yonekawa T, Watanabe K, Amino N, et al. Loop-mediated isothermal amplification of DNA. Nucleic Acids Res. 2000;28(12):e63–e. doi: 10.1093/nar/28.12.e63.

28. Mori Y, Kanda H, Notomi T. Loop-mediated isothermal amplification (LAMP): recent progress in research and development. J Infect Chemother. 2013;19(3):404–11. doi: 10.1007/s10156-013-0590-0.

29. Almassian DR, Cockrell LM, Nelson WM. Portable nucleic acid thermocyclers. Chem Soc Rev. 2013;42(22):8769–98. doi: 10.1039/c3cs60144g.

30. Peters IR, Helps CR, Batt RM, Day MJ, Hall EJ. Quantitative real-time RT-PCR measurement of mRNA encoding α-chain, pIgR and J-chain from canine duodenal mucosa. J Immunol Methods. 2003;275(1):213–22. doi: 10.1016/S0022-1759(03)00056-5.

31. Fang W, Wang J, Liu J, Xu C, Cai W, Luo D. PCR assay for the cell-free copro-DNA detection of Angiostrongylus cantonensis in rat faeces. Vet Parasitol. 2012;183(3):299–304. doi: doi.org/10.1016/j.vetpar.2011.07.026.

32. Mallaiyaraj Mahalingam JT, Calvani NED, Lee R, Malik R, Šlapeta J. Using cerebrospinal fluid to confirm Angiostrongylus cantonensis as the cause of canine neuroangiostrongyliasis in Australia where A. cantonensis and Angiostrongylus mackerrasae co-exist. Curr Res Parasitol Vector Borne Dis. 2021;1:100033. doi: 10.1016/j.crpvbd.2021.100033.

33. Valentyne H, Spratt DM, Aghazadeh M, Jones MK, Šlapeta J. The mitochondrial genome of Angiostrongylus mackerrasae is distinct from A. cantonensis and A. malaysiensis. Parasitology. 2020;147(6):681–8. doi: 10.1017/S0031182020000232.

34. Jia B, Li X, Liu W, Lu C, Lu X, Ma L, et al. GLAPD: whole genome based LAMP primer design for a set of target genomes. Front Microbiol. 2019;10:2860. doi: 10.3389/fmicb.2019.02860.

35. Landis JR, Koch GG. The measurement of observer agreement for categorical data. Biometrics. 1977;33(1):159–74. doi: 10.2307/2529310.

36. Giavarina D. Understanding Bland Altman analysis. Biochem Med (Zagreb). 2015;25(2):141–51. doi: 10.11613/bm.2015.015.

37. Spratt DM. Species of Angiostrongylus (Nematoda: Metastrongyloidea) in wildlife: A review. Int J Parasitol Parasites Wildl. 2015;4(2):178–89. doi: 10.1016/j.ijppaw.2015.02.006.

38. Prociv P, Carlisle MS. The spread of Angiostrongylus cantonensis in Australia. Southeast Asian J Trop Med. 2001;32 Suppl 2:126–8.

39. Mackie J, Lacasse C, Spratt D. Patent Angiostrongylus mackerrasae infection in a black flying fox (Pteropus alecto). Aust Vet J. 2013;91(9):366–7. doi: 10.1111/avj.12082.

40. Seebeck J, Menkhorst P. Status and conservation of the rodents of Victoria. Wildl Res. 2000;27(4):357–69. doi: 10.1071/WR97055.

41. Peakall R, Ebert D, Cunningham R, Lindenmayer D. Mark–recapture by genetic tagging reveals restricted movements by bush rats (Rattus fuscipes) in a fragmented landscape. J Zool. 2006;268(2):207–16. doi: 10.1111/j.1469-7998.2005.00011.x.

42. Fox BJ, Monamy V. A review of habitat selection by the swamp rat, Rattus lutreolus (Rodentia: Muridae). Austral Ecol. 2007;32(8):837–49. doi: 10.1111/j.1442-9993.2007.01849.x.

43. Dusitsittipon S, Criscione CD, Morand S, Komalamisra C, Thaenkham U. Hurdles in the evolutionary epidemiology of Angiostrongylus cantonensis: Pseudogenes, incongruence between taxonomy and DNA sequence variants, and cryptic lineages. Evol Appl. 2018;11(8):1257–69. doi: 10.1111/eva.12621.

44. Červená B, Modrý D, Fecková B, Hrazdilová K, Foronda P, Alonso AM, et al. Low diversity of Angiostrongylus cantonensis complete mitochondrial DNA sequences from Australia, Hawaii, French Polynesia and the Canary Islands revealed using whole genome next-generation sequencing. Parasit Vectors. 2019;12:241. doi: 10.1186/s13071-019-3491-y.

45. Tokiwa T, Harunari T, Tanikawa T, Komatsu N, Koizumi N, Tung K-C, et al. Phylogenetic relationships of rat lungworm, Angiostrongylus cantonensis, isolated from different geographical regions revealed widespread multiple lineages. Parasitol Int. 2012;61(3):431–6. doi: 10.1016/j.parint.2012.02.005.

46. Monte TCC, Simões RO, Oliveira APM, Novaes CF, Thiengo SC, Silva AJ, et al. Phylogenetic relationship of the Brazilian isolates of the rat lungworm Angiostrongylus cantonensis (Nematoda: Metastrongylidae) employing mitochondrial COI gene sequence data. Parasit Vectors. 2012;5(1):248. doi: 10.1186/1756-3305-5-248.

47. Monte TC, Gentile R, Garcia J, Mota E, Santos JN, Maldonado Júnior A. Brazilian Angiostrongylus cantonensis haplotypes, ac8 and ac9, have two different biological and morphological profiles. Mem Inst Oswaldo Cruz. 2014;109:1057–63. doi: 10.1590/0074-0276130378

48. Lee J-D, Chung L-Y, Wang L-C, Lin R-J, Wang J-J, Tu H-P, et al. Sequence analysis in partial genes of five isolates of Angiostrongylus cantonensis from Taiwan and biological comparison in infectivity and pathogenicity between two strains. Acta Trop. 2014;133:26–34. doi: 10.1016/j.actatropica.2014.01.010.

49. Jacob J, Steel A, Lin Z, Berger F, Zöeller K, Jarvi S. Clinical efficacy and safety of albendazole and other benzimidazole anthelmintics for rat lungworm disease (neuroangiostrongyliasis): A systematic analysis of clinical reports and animal studies. Clin Infect Dis. 2021;74(7):1293–302. doi: 10.1093/cid/ciab730.

50. Prociv P, Turner M. Neuroangiostrongyliasis: The “subarachnoid phase” and its implications for anthelminthic therapy. Am J Trop Med. 2018;98(2):353–9. doi: 10.4269/ajtmh.17-0206.

